# Determinants of visual ambiguity resolution

**DOI:** 10.1101/2025.05.28.656283

**Authors:** Juan Linde-Domingo, Javier Ortiz-Tudela, Johannah Völler, Martin N. Hebart, Carlos González-García

**Author notes:** These authors contributed equally (Juan Linde-Domingo, Javier Ortiz-Tudela). Correspondence to Juan Linde-Domingo or Carlos González-García.

## Abstract

Visual inputs during natural perception are highly ambiguous: objects are frequently occluded, lighting conditions vary, and object identification depends significantly on prior experiences. However, why do certain images remain unidentifiable while others can be recognized immediately, and what visual features drive subjective clarification? To address these critical questions, we developed a unique dataset of 1,854 ambiguous images and collected more than 100,000 ratings (from a total of 947 participants) evaluating their identifiability before and after seeing undistorted versions of the images. Relating the representations of a brain-inspired neural network model in response to our images with human ratings, we show that subjective identification depends largely on the extent to which higher-level visual features from the original images are preserved in their ambiguous counterparts. In line with these results, an image-level regression analysis showed that the subjective identification of ambiguous images was best explained by high-level visual dimensions. Notably, the predominance of higher-level features over lower-level ones softens after participants disambiguate the images, suggesting that the visual system flexibly shifts between top-down guessing to bottom-up matching after disambiguation. Moreover, we found that the process of ambiguity resolution was accompanied by a notable decrease in semantic distance and a greater consistency in object naming among participants. However, the relationship between information gained after disambiguation and subjective identification was non-linear, indicating that acquiring more information does not necessarily enhance subjective clarity. Instead, we observed a U-shaped relationship, suggesting that subjective identification improves when the acquired information either strongly matches or mismatches prior predictions. Collectively, these findings advance our understanding on how we resolve ambiguity and extract meaning from incomplete visual information.

## Introduction

The natural world is inherently ambiguous, yet humans can in most cases make sense of what they see. This remarkable feature of human perception is a core principle of the predictive processing framework, which casts perception as a purely inferential process of hypothesis testing ^1–4^. Specifically, to resolve ambiguity, this framework proposes that the brain generates predictions about the most probable causes underlying the sensory input from real-world experience. These initial predictions are then compared to feedforward sensory input, and any mismatch – or prediction error – drives an iterative refinement process. Through repeated prediction error minimization, perception stabilizes, and ambiguity is resolved ^2,5^.

Despite the central role of ambiguity in predictive processing, experimental research has rarely tested how iterative prediction error minimization unfolds when encountering truly ambiguous stimuli. Instead, studies on visual object recognition typically rely on clear and unambiguous images ^6–8^ or more complex, “challenging” natural images ^9^. These studies robustly demonstrate that top-down prior knowledge influences perception, even at early processing stages (i.e., areas V1, V2 of the primate visual cortex ^5,10^). Despite the significance of these studies, we still lack an understanding of the cognitive and input-related determinants of visual ambiguity and how we can resolve it. This is the case primarily because it remains difficult to experimentally capture the ambiguity inherent in real-world perception, which arises mostly from two sources: (1) the biological limitations of the sensory system, such as structural limitations or inherent neural noise, and (2) a sensory input that is often incomplete or unclear. While the former can be studied even with clear stimuli, the latter requires the use of impoverished materials mimicking real-world scenarios ^11^.

Importantly, in real-world perception, ambiguity is rarely passively endured; instead, we actively seek additional information to resolve it ^12^. For example, imagine walking down a road and noticing a shadowy shape in the distance. This shape is ambiguous not only because of the limitations of our sensory organs, but also because it happens to be partially occluded by a tree, which, additionally, casts shadows over some part of the shape, thus altering the input that actually reaches the eyes. Your brain, based on prior experience, predicts it is a road sign. However, the incomplete visual information makes it difficult to verify your hypothesis. To resolve this ambiguity, you might change your viewing angle, move closer, or squint – actively gathering new inputs to refine your prediction and minimize the error. Several questions arise at this point: what makes the sign ambiguous in the first place? What information was missing or distorted to prevent identification? ^11^. Also, what is the effect of this information-seeking behaviour on your perception?

Here, we build upon these ideas to (i) determine why certain stimuli induce ambiguity and others do not, and (ii) reveal the relationship between gaining new information and subjective disambiguation. To address these research questions, we use ambiguous black-and-white Mooney images ^13^, which are difficult to identify without additional information, but become effortlessly identifiable afterwards. These stimuli allow for real-time investigation of ambiguity resolution in a controlled setting, allowing for a direct test of how perception evolves when new information is gained ^14–17^, mirroring natural perceptual processes.

Answering these questions requires a large amount of data but, to date, no exhaustive collection of Mooney images is publicly available. Therefore, we generated a large-scale, openly available dataset of Mooney images of 1,854 objects. To do so, we created the Mooney version of the images from the THINGSplus database ^18^ and paired it with over 100,000 behavioural ratings from more than 1,000 participants. We first verified that the curated images worked as expected, in that they were poorly identified unless an external, unambiguous input was provided. Next, we examined what makes stimuli ambiguous in the first place by deriving feature representations from text-based embeddings and human behaviour, in combination with deep neural networks (DNNs) that emulate hierarchical processing in the ventral stream ^19,20^. We then determined what information helps to resolve ambiguity and how this newly gained information shapes subsequent subjective identification. To this end, we first explored which specific combination of visual features relates to subjective identification before and after disambiguation. Then, we implemented two metrics that identify the type of semantic information that can be learned during disambiguation and their relationship with subjective identification.

Based on this framework, we hypothesised that the successful resolution of visual ambiguity requires a dynamic interplay between different levels of visual representation. Specifically, we expected subjective identification to depend initially on the preservation of high-level information, to then shift toward matching these predictions with specific low-level features. To anticipate our results, we observed that ambiguous stimuli are prevented from subjective identification when higher-level, rather than low-level visual features are distorted. However, after disambiguation, lower-level features become more related to subjective identification while higher-level ones play a reduced role. Moreover, disambiguation leads to significant changes in the semantic representations induced by Mooney images, reducing their distance in semantic space to their original, undistorted counterpart while also decreasing uncertainty. Finally, our results reveal that the relationship between information gain and later subjective identification is not strictly linear. Instead, we observe a U-shaped pattern, suggesting that knowledge acquisition does not directly translate into improvements in subjective identification. Rather, later identification improves whenever the acquired information either strongly confirms or violates previous predictions.

## Methods

### Participants

A total of 1065 participants were recruited online via Prolific. Participants who did not finish the task or presented mean reaction times longer than 5 s in the identification task were excluded, remaining 1002 of them (384 identified as female, 608 male, 7 answered “other”, and 3 participants preferred not to answer; mean age 28.67, SD ±4.60; 831 from the United Kingdom, 153 from the United States of America, and 13 from other countries; data on ethnicity was not collected). The eligibility criteria were that participants had to be between 18 and 35 years old and have English as a first language. Additionally, only participants with an approval rate between 95-100% and more than 10 previous submissions in other experiments in Prolific were eligible. No data on socioeconomic status, race, or ethnicity was collected. Regarding exclusion criteria, participants from previous pilot experiments with the same images, as well as participants using smartphones or tablets, were not eligible. For the remaining participants, we applied the 1.5 interquartile range (IQR) rule in the identification task for unambiguous images to exclude outliers (55 outliers in the lower bound). After this, 947 participants remained for further analyses. All participants were reimbursed at a rate of £6.00 per hour for completing the task (∼ 20 minutes); participation was entirely voluntary, and all participants provided informed consent. The experiment was approved by the Ethics Committee of the University of Granada.

### Stimuli, task, and procedure

A total of 1854 greyscale-transformed images from the THINGSplus object database ^18^ were used in the experiment. Three experimenters, who had been previously trained, transformed all greyscale images into Mooney images by manually adjusting a Gaussian filter and selecting an intensity threshold for each item individually. This threshold operated so that pixels above it are set to white and pixels below are set to black, rendering two-tone images.

In the main task, each trial started with a fixation cross (1 s) in the centre of the screen, followed by either a Mooney or a greyscale unambiguous image. Participants were asked to respond as quickly as possible whether they identified or not the object on the displayed image by pressing the “E” or “I” key respectively (response mappings were counterbalanced across participants). After response or 5 s, participants were asked to type the name of the object using the keyboard. This naming task was intended to verify participants’ identification. When participants started typing, they were prompted with a drop-down menu that contained a series of words matching the typed letters. Participants were instructed to select the desired word by clicking on it. These words included the THINGS image names plus their WordNet synonyms, 3434 words in total. The next trial started after choosing the name of the object or after a maximum of 20 s. The Mooney version of each image was presented in two different trials (from now on, pre– and post-disambiguation), always interleaved by a trial where its unambiguous, greyscale version was displayed (disambiguation trial). The maximum distance between a Mooney image and its unambiguous version was 4 trials. Due to design constraints reasons, the Mooney and unambiguous version of the same image were presented sequentially in one case per participant. The experiment lasted approximately 20 minutes.

Participants were collected following a batch fashion. To sample the entire image space, the 1854 images were randomly divided in 46 subgroups of 40 images each. Then, we proceeded to collect data from 3 batches (∼150 subjects, to account for dropouts). Since participants were randomly allocated to one of the 47 subgroups, some images were oversampled while others were not presented at all. To account for this, every 3 batches, we collected an additional fine-tuned batch that contained those subgroups of images that were not presented initially. Additionally, due to a technical issue, participants from the first 9 batches had some words (394 out of 3434) missing from the drop-down menu of the naming task. To avoid any potential influence of this issue, we removed these trials from these participants. Moreover, to compensate for these missing trials, we ran one additional batch of 100 participants that were presented 40 of the 394 images each, before continuing with the 10th batch of participants. On average, each participant rated 34.10 images (±5.28; range = [10 – 40]). Across all participants, images rated by participants a number of times below 2 SD of the mean number of ratings were excluded from further analysis (40 out of a total of 1854 images). After applying this exclusion criteria, each of the remaining 1814 images was rated an average of 17.80 times (±3.51; range = [11 – 33]). An additional five images had to be excluded from semantic distance analyses because their semantic embedding was not available in the original THINGSplus object database ^18^. DNN analyses were conducted on this reduced set. One additional image (“switch”) was not included due to a preprocessing issue. The study was not preregistered.

### Data preprocessing

After exclusion of participants and images (see above), we processed the provided verbal labels for the remaining trials to check for the use of synonyms (considering those included in WordNet for each of our true labels) and computed semantic distance as the cosine dissimilarity between the embeddings of the provided and true label. Embeddings were retrieved from the THINGS metadata. Specifically, we used a word2vec implementation of sense vector (developed to distinguish between different meanings of words) augmented to account for missing entries. We then computed semantic entropy to quantify response diversity for each image under different conditions. For each image-condition pair, response probabilities were derived by normalizing the frequency of each unique verbal descriptor. Entropy was then calculated applying Shannon’s formula to the normalized frequencies. No further preprocessing was done except for reaction time analysis, where trials with responses below 200 milliseconds or above 5 seconds were excluded.

### Image-level correlation estimation

To obtain an image-level correlation between their preservation index and subjective identification values, we implemented a bootstrapped random sampling approach inspired by approaches to obtain local correlations ^39^. This method calculates the correlation coefficients for subsets of data associated with each image, while introducing randomness to enhance robustness. Image-level correlations were calculated per image, using subsets of other images of a given size (n = 3). These subsets were defined randomly without replacement (even in other iterations of the bootstrapping approach), ensuring variability in the selection across repetitions. Pearson correlation coefficient was calculated between the two variables for each randomly selected subset. To obtain a more stable estimate, the random selection and local correlation were repeated 400 times (bootstrapping), keeping in mind the limitation of defining each dataset without replacement. For each image, this resulted in 400 local correlation values which were then averaged to calculate the image-level correlation.

A null correlation baseline was generated following the same approach but shuffling the two variables independently within each subset. This preserved the statistical properties of the data while removing any inherent relationship between variables. The same bootstrapping and aggregation procedures were applied to the shuffled data.

### Regression on dimensional embedding

Out of the total 49 available object space dimensions for each image obtained in the THING database ^27^, a subset of 36 were labeled as mostly semantic or mostly visual based on previous study using a similar approach ^27^. Each dimensional embedding was treated as a predictor in a regression model for subjective identification in post, enabling the decomposition of the total variance explained (\( R^2 \)) into components attributable to semantic features, visual features, and their shared interactions.

To contextualize the model’s performance within the limits of the data’s inherent predictability, we computed a noise ceiling representing the maximum explainable variance. This was computed using a split-half reliability approach repeated 1,000 times to ensure robustness. In each iteration, participants were randomly split into two non-overlapping groups, and average subjective identification scores were computed for each image within each group. The Pearson correlation between the two groups’ image-level scores was calculated and corrected using the Spearman-Brown prediction formula to estimate the reliability of the full dataset. The average Spearman-Brown–corrected value across iterations was used as the noise ceiling. Model performance was then expressed as a proportion of this ceiling, indicating how much of the explainable variance was accounted for by the dimensions.

In addition to this global measure, variance partitioning was performed to disentangle the unique contributions of semantic and visual dimensions, as well as their shared variance. Variance partitioning involved fitting regression models for semantic dimensions alone, visual dimensions alone, and both combined. From these models, we computed the unique variance explained by semantic dimensions and the unique variance explained by visual dimensions. These components were normalized relative to the model-explained variance to obtain a reliable estimate of the relative contribution of each feature type under the constraints of the data’s inherent noise.

Given the large sample sizes and the robustness of parametric tests to moderate non-normality, normality was assessed via residual inspection rather than formal tests ^40^.

## Results

### Large stimulus set validates established disambiguation effects

A total of 1,065 participants (final n after exclusions = 947; see Methods section) performed an online visual identification task (Fig. 1b). On each trial, an image was presented on the screen. The images were taken from the THINGSplus database ^18^ and reflected a broad set of 1,854 images of natural objects. The images were either presented as an unambiguous, greyscale version of these objects, or as an ambiguous two-tone Mooney image that was created by adding Gaussian blur and binarization (Fig. 1a, see Methods), making it a total of 3,708 images, 3,628 of which were included in further analyses (see Methods for image exclusion details). Participants were first asked to respond whether they could identify the main object on the image. Subsequently, to assess the extent to which participants’ subjective identification matched the true content of the images, they were asked to provide a verbal label or to guess a label if they were unsure about the correct one. The Mooney version of each image was presented in two different trials, always interleaved by a trial where its unambiguous version was displayed (maximum distance between a Mooney and its unambiguous greyscale version = 4 trials, see Methods), allowing us to experimentally induce visual disambiguation. In the following, we will refer to these as pre-disambiguation trials (Mooney images), disambiguation trials (unambiguous, greyscale images), and post-disambiguation trials (Mooney images). This nomenclature reflects our experimental approach, manipulating the timing of ambiguity resolution, and follows the terminology used in previous research ^15,21^. Participants were not explicitly informed about different trial types. Each of the 3,628 images was rated by participants an average of 17.8 times (±3.51; range = [11-33]).

**Fig. 1.**
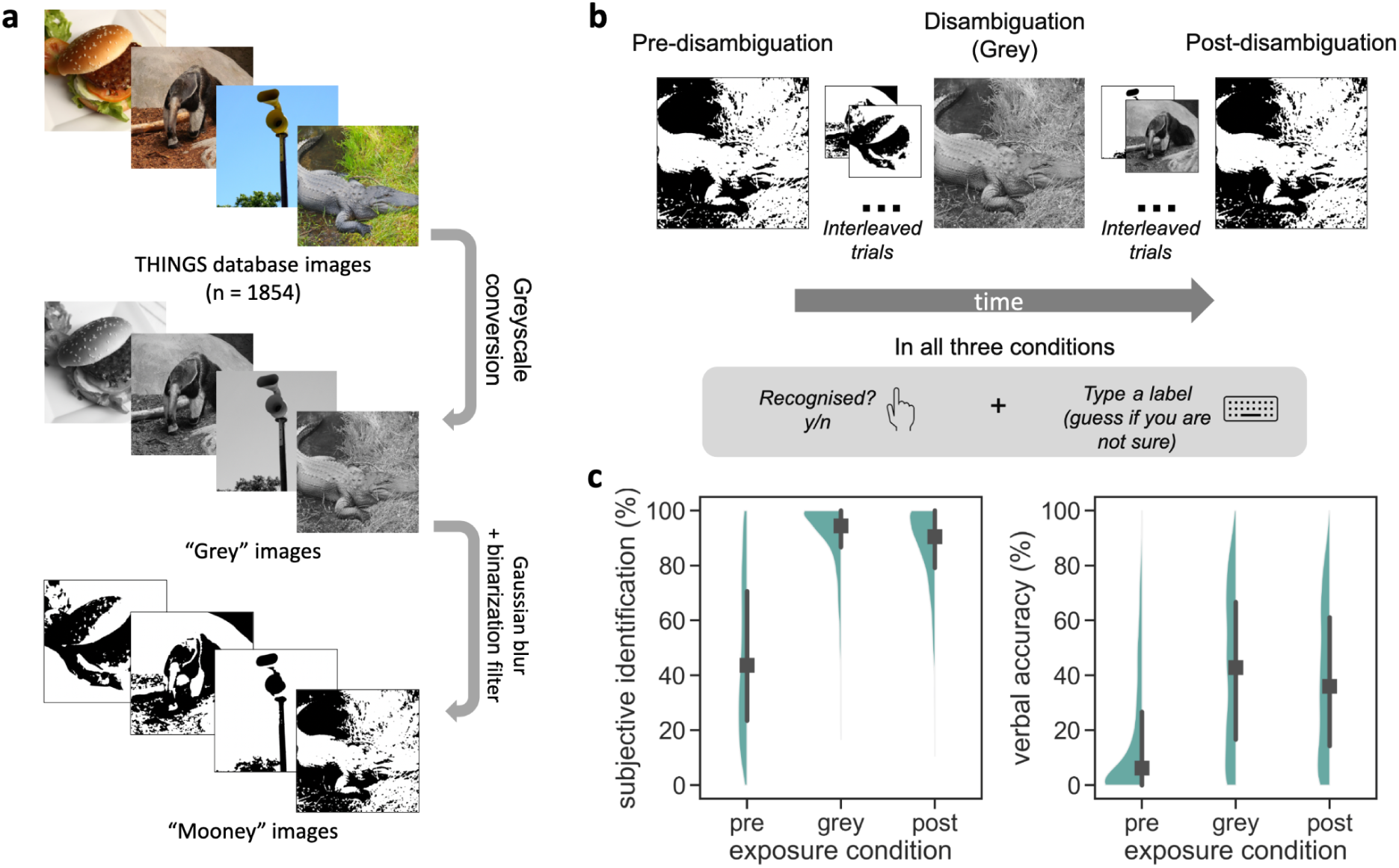
Experimental design and behavioural performance. **a**, A total of 1,854 images obtained from the THINGS database were transformed into Mooney images. This was achieved converting these images into greyscale and applying a Gaussian blur and binarization filter (see Methods section). **b**, 1,065 participants were presented with images in three conditions: pre-disambiguation (initial exposure to Mooney images), disambiguation (presenting the unambiguous greyscale image), and post-disambiguation (re-exposure to Mooney images after seeing their unambiguous version). In all conditions, participants responded whether they identified the image (yes/no) and typed a label. **c**, Subjective identification rates (left) and verbal accuracy (right) across all three conditions (n = 947 participants per condition). Black squares represent medians, black lines denote the interquartile range, and green shaded areas indicate the data distribution. Unambiguous images in the figure are extracted from THINGSplus ^18^ and are copyright-free. Mooney images were generated based on this material and are copyright-free as well. Behavioural analyses are based on n = 947 participants after exclusions and n = 1,814 images (each rated on average 17.8 times, range 11–33).

To evaluate whether the manipulation worked as intended, we first investigated whether subjective identification was affected by exposure conditions. As expected ^14,15,22–24^, a repeated-measures ANOVA revealed that subjective identification increased significantly (F_(2, 3626)_ = 4101.09, p < 0.001, η²_p_= 0.69, 95% CI [0.68, 0.71]; Fig. 1C, left panel) after exposure to the unambiguous version of the image (M = 85.9%, SD = 15.1%), compared to pre-disambiguation trials (M = 47%, SD = 27.9%). Similarly, reaction times were significantly faster after (M = 1692 ms, SD = 366) compared to pre-disambiguation (M = 2537 ms, SD = 410; F_(2, 3626)_ = 2902, p < 0.001, ηp2 = 0.62, 95% CI [0.60, 0.63]). To assess correct identification, we measured performance in the subsequent naming task, considering both the provided label for each image and the true label plus a list of predefined synonyms as correct responses. Participants’ precision in naming increased significantly after disambiguation trials (pre-disambiguation, M = 16.7%, SD = 22.7%; post-disambiguation, M = 38.5%, SD = 26.9%; F_(2, 3626)_ = 2092, p < 0.001, ηp2 = 0.54, 95% CI [0.52, 0.55]; see Fig. 1C, right panel). Of the 85.9% (SD = 15.1%) trials where participants indicated identifying the object post-disambiguation, they typed the correct label (semantic distance = 0, including synonyms; see below) 47.8% (SD = 17.92%) of them. Altogether, these analyses confirm that the material used robustly produced the expected behavioural enhancement in subjective and objective measures in identification of ambiguous images after disambiguation, in line with previous results ^14,15,24^.

### Feature preservation and subjective identification

Resolving visual ambiguity likely depends on a combination of the visual attributes of the stimuli at different levels of complexity, ranging from lower-level features (e.g., line orientations) to higher-level ones (e.g., object identity) ^25^. To explore the relative contribution of these types of features to subjective identification, we used a computational approach inspired by the hierarchical architecture of the human ventral visual stream. This analysis involved two main steps: feature extraction across hierarchical layers of a neural network model, and estimation of the degree to which feature representations of the unambiguous images are maintained in the Mooney images via a similarity-based assessment (Fig. 2a).

**Fig. 2.**
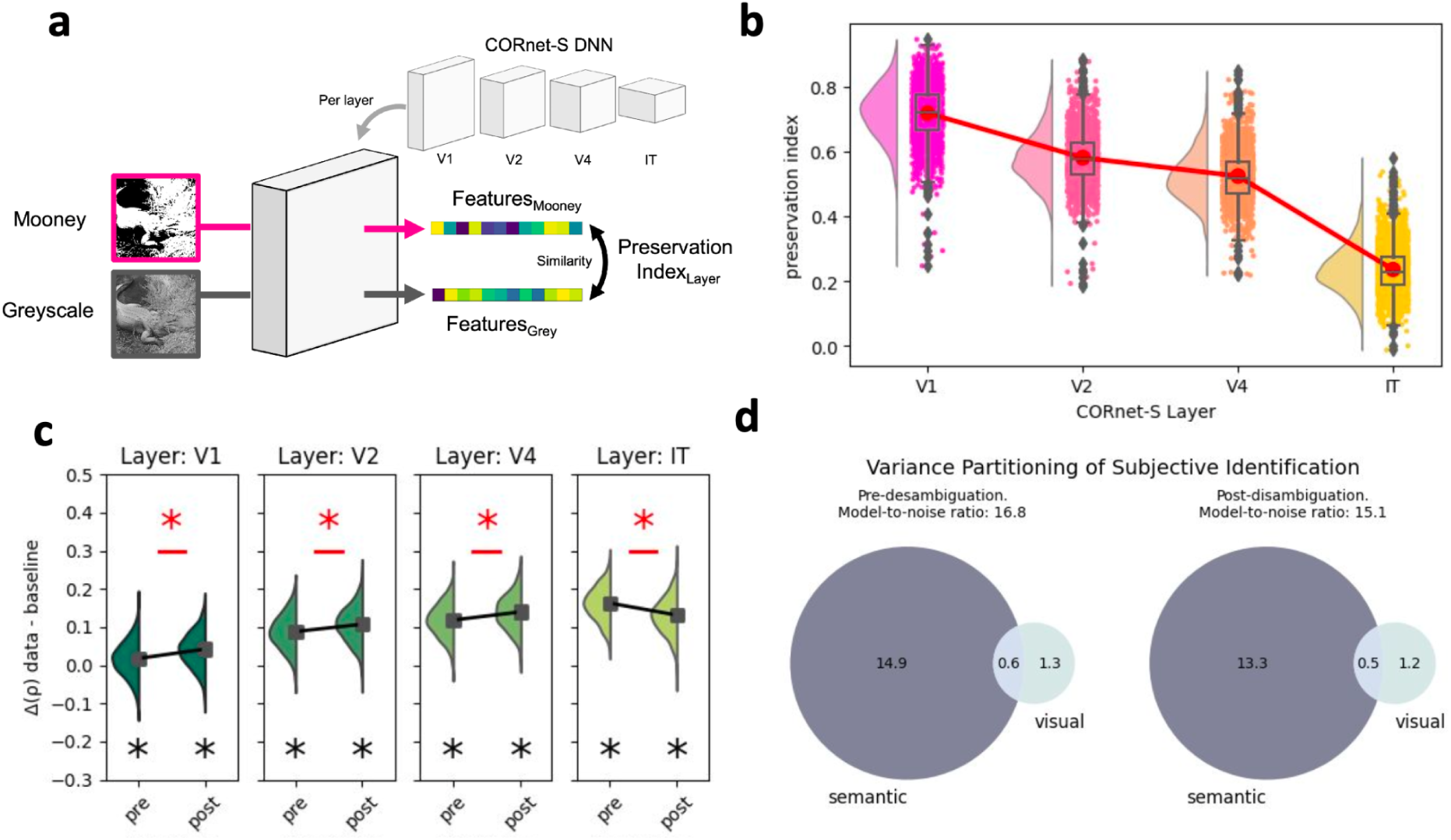
Visual and verbal stimulus features and their link to subjective identification. **a**, Mooney and unambiguous greyscale versions of the same image were processed through CORnet-S, a DNN that mimics the hierarchical architecture of the ventral visual stream. For each layer, feature representations were extracted, and the similarity (Pearson) between Mooney and unambiguous features was calculated as the preservation index. **b**, The preservation index decreased progressively from early visual layers (V1) to higher-level layers (IT), indicating that higher-level representations are more impaired by the Mooney transformation. **c**, Differences in preservation index and subjective identification correlations across DNN layers and exposure conditions (pre– and post-disambiguation) are shown. A significant increase in correlation was observed in V1 and V2 after disambiguation (red asterisks), while a decrease occurred in IT, suggesting that lower-level features play a stronger relationship after disambiguation. d, Venn diagrams illustrate the contribution of verbal semantic and visual dimensions to variance in subjective identification pre– and post-disambiguation. Preservation index and preservation–identification association analyses were conducted at the image level (n = 1,808 images, see Methods). Regression/variance partitioning analyses were conducted at the image level (n = 1,814 images). Subjective identification values were derived from n = 947 participants.

First, unambiguous greyscale and Mooney versions of the same image were processed through CORnet-S ^20^. This deep convolutional neural network is a computational model that emulates hierarchical processing in the primate ventral stream with separate modules for V1, V2, V4, and IT and has been shown to yield a good correspondence of individual layers to different cortical processing stages ^20^. For each input image, we extracted the feature representations at each module within the network using the Net2Brain toolbox ^26^. In the second step, we quantified the similarity between the features extracted from the unambiguous grayscale and Mooney versions of each image. This similarity, measured as a Pearson correlation between the two feature sets at each layer, yielded a preservation index for each image at each level (Fig. 2a). The preservation index reflects the degree to which properties of the original image were maintained in its ambiguous counterpart on a specific level of the processing hierarchy of the visual system. In other words, this index provides insights into how much of the original image’s representational content survives the abstraction process of creating Mooney images. Across all layers, the preservation index was significantly greater than zero. Specifically, preservation was highest in V1 (M = 0.719, SD = 0.084, 95% CI [0.715, 0.723]), *t*(1807) = 364.341, *p* < 0.001, Cohen’s *d* = 8.569, and decreased in V2 (M = 0.581, SD = 0.082, 95% CI [0.578, 0.585]), *t*(1807) = 302.949, *p* < .001, Cohen’s *d* = 7.125, V4 (M = 0.525, SD = 0.080, 95% CI [0.521, 0.528]), *t*(1807) = 279.331, *p* < 0.001, Cohen’s *d* = 6.569, and IT (M = 0.236, SD = 0.072, 95% CI [0.233, 0.240]), *t*(1807) = 140.128, *p* < 0.001, Cohen’s *d* = 3.296. These results indicate that Mooney images consistently retained features of their unambiguous counterparts across all processing stages, although the degree of preservation decreased along the hierarchy (Fig. 2b). Accordingly, a one-way ANOVA revealed significant differences in preservation across layers, *F*(3, 7228) = 11840.99, *p* < 0.001, η²p = .831, 95% CI [.612, .630], indicating that original features of the unambiguous image were particularly impaired in higher-level layers after Mooney transformation. To better assess whether the preservation index pattern across layers was driven by our image manipulation, we conducted control analyses using alternative high-pass and low-pass filtered transformations (see Supplementary Materials, Feature preservation for non-Money image transformations section). The results showed that the high-pass transformation produced a distinct preservation pattern, with equivalent preservation in early and late layers. In contrast, the low-pass transformation yielded a preservation profile similar to that of the Mooney transformation, with reduced preservation in higher-level layers. This result is consistent with the construction process of the Mooney images, which included a low-pass Gaussian filter as one of its steps.

We then explored the impact of this particular pattern of feature retention on behavior. In particular, we examined 1) the extent to which the degree of preservation of individual images correlated with their subjective identification during Mooney image presentation, and 2) whether such identification was more strongly associated with the preservation of lower-level features or higher-level information. To do so, for each image, we calculated the image-level correlation (see Methods) between the preservation index and subjective identification. This correlation was computed for each DNN layer and exposure condition (pre– and post-disambiguation).

To assess whether these associations were reliably above baseline, we tested them against zero using one-sided paired *t*-tests (Bonferroni corrected). In the pre-disambiguation condition, V1 showed a modest but significant positive association (*t*(1807) = 11.202, *p* < .001, Cohen’s *d* = 0.263, 95% CI [0.010, 0.014]). Stronger positive associations were observed in V2 (*t*(1807) = 81.174, *p* < 0.001, *d* = 1.909, 95% CI [0.087, 0.091]), V4 (*t*(1807) = 108.983, *p* < 0.001, *d* = 2.563, 95% CI [0.116, 0.120]), and IT (*t*(1807) = 149.488, *p* < .001, *d* = 3.516, 95% CI [0.156, 0.160]). A similar positive pattern to the pre-disambiguation condition was observed in the post-disambiguation condition (V1: *t*(1807) = 36.443, *p* < .001, *d* = 0.857, 95% CI [0.039, 0.043]; V2: *t*(1807) = 100.589, *p* < 0.001, *d* = 2.366, 95% CI [0.109, 0.113]; V4: *t*(1807) = 130.016, *p* < 0.001, *d* = 3.058, 95% CI [0.140, 0.145]; IT: *t*(1807) = 120.966, *p* < 0.001, *d* = 2.845, 95% CI [0.130, 0.135]).These results indicate that higher preservation of image features was consistently associated with greater subjective identification across all layers and conditions (Fig. 2c).

Furthermore, a two-way repeated-measures ANOVA with factors of layer and exposure condition revealed a significant interaction between layer and condition, F(6, 10842) = 1367.948, p < 0.001, ηp² = .431, 95% CI [.107, .117], together with significant main effects of layer, F(3, 5421) = 10888.548, p < 0.001, ηp² = .858, 95% CI [.658, .677], and exposure condition, F(2, 3614) = 56216.173, p < 0.001, ηp² = .969, 95% CI [.937, .942]. As expected, these main effects reveal that the relation between subjective identification and preservation 1) becomes stronger in higher-order layers, indicating an overall greater reliance on higher-level features for subjective identification; and 2) is generally higher after disambiguation. These results, first, confirm that CORnet-S-based preservation indices capture significant variation in human representational content. Importantly, they provide solid grounds for further analyses of the interaction effect suggesting that this general pattern is reversed in higher-order layers.

More specifically, planned paired *t*-test comparisons revealed a significant increase in the association between preservation and subjective identification after disambiguation in early and intermediate layers, including V1, *t*(1807) = −18.472, *p* < 0.001, Cohen’s *d* = −0.434, 95% CI [−0.032, −0.026], V2, *t*(1807) = −14.271, *p* < 0.001, *d* = −0.336, 95% CI [−0.026, −0.019], and V4, *t*(1807) = −15.950, *p* < 0.001, *d* = −0.375, 95% CI [−0.028, −0.022] (all Bonferroni corrected). In contrast, a significant decrease was observed in the IT layer, *t*(1807) = 17.058, *p* < 0.001, *d* = 0.401, 95% CI [0.023, 0.029] (Bonferroni corrected). These results suggest that the preservation of lower-level features (as captured in V1, V2 and V4) increases its relative effect on subjective identification after disambiguation, compared to higher-level features (e.g., IT), whose influence diminishes once the unambiguous greyscale image is presented.

Additionally, we explored the role of other mnemonic factors that could contribute to later identification. First, we obtained memorability scores for all unambiguous images from a previous study ^27^ and computed Pearson correlations between these values and our behavioural measures of subjective identification and semantic distance across conditions. This analysis tested whether image memorability could account for differences in subjective identification, semantic distance, or information gain during disambiguation. Overall, we found a series of weak correlations, indicating that memorability is not specially connected to subjective recognition or semantic distance in ambiguous conditions. Specifically, although more memorable images tended to be recognised better in the unambiguous greyscale condition (*r*(1845) = 0.112, *p* < 0.001, 95% CI [0.067, 0.157]), this positive relationship disappeared for ambiguous Mooney images.

Second, we examined whether identification in post-disambiguation trials could be driven by repeated exposure to items from the same category, which might bias responses. For each participant, we computed the correlation between the number of images shown per category and their average subjective recognition and semantic distance and then tested whether the mean correlation across participants differed from zero. This analysis provided no evidence for such an effect. The mean group correlation between category frequency and subjective recognition was not significant in the positive direction and, if anything, negative (mean *r* = −0.118, 95% CI [−0.140, −0.097]; *t*(839) = −10.77, p = 1.00, one-tailed; Cohen’s *d* = −0.37, 95% CI [−0.44, −0.30]).. Similarly, there was no statistically significant relationship in the negative direction between category frequency and semantic distance and, if anything, the association was slightly positive (mean *r* = 0.033, 95% CI [0.013, 0.053]; *t*(940) = 3.20, p = 0.999, one-tailed; Cohen’s *d* = 0.10, 95% CI [0.04, 0.17]).

### Contribution of visual dimensions to identification

Building on these findings, we next determined which specific combination of visual dimensions enables successful subjective identification (pre– and post-disambiguation) using the metadata of the THINGS database ^18,28,29^. The THINGS database provides a 66-dimensional behavioural similarity embedding for the 1,854 object concepts used in this study, constructed based on human similarity judgments ^29,30^. These dimensions have been shown to capture visual (e.g., round, fine-grained) and semantic (e.g., animal-related, valuable) properties of objects. In our study, three independent raters classified this original set of dimensions as either visual, semantic or mixed. Then, we discarded the ones labeled as mixed and all color-related dimensions (e.g., “sand-colored”) as our stimuli were greyscale images. This left us with 47 dimensions. We next applied a multiple regression approach which allowed us to simultaneously evaluate the unique and shared predictive power of these dimensions. To contextualize this result, we calculated a noise ceiling over 1,000 participant splits using Spearman-Brown corrected split-half reliability, representing the maximum explainable variance, given the inherent noise in the data.

In the pre-disambiguation condition, the regression model explained a total variance (R^2^) of 0.153, *F*(47, 1766) = 6.48, *p* < 0.001, Cohen’s f^2^ = 0.18, in subjective identification ratings. The average noise ceiling, representing the maximum explainable variance, was 0.925, 95% CI [.919, .930], resulting in a model-to-ceiling ratio of 0.18. This indicates that 18% of the explainable variance was captured by the model, suggesting that while the model may leverage meaningful predictive information from the dimensional embeddings, a substantial portion of the variance remains unexplained, likely reflecting other sources of variability (e.g., trial-by-trial changes within-participants). Variance partitioning revealed that 62.4% of the total explained variance was uniquely attributable to semantic dimensions, 21.8% to visual dimensions, and the remaining portion to shared contributions between the two. These results highlight the dominant role of semantic information in shaping identification judgments prior to disambiguation, with visual dimensions providing minimal additional predictive value.

In the post-disambiguation condition, the regression model explained a total variance (R^2^) of 0.099, *F*(47, 1766) = 3.88, *p* < 0.001, Cohen’s f^2^ = 0.11, in subjective identification ratings. The average noise ceiling was 0.849, 95% CI [.834, .861], with a model-to-ceiling ratio of 0.137, indicating that the model captured approximately 14% of the maximum explainable variance. Despite this decrease in overall predictive performance, the relative contributions of semantic and visual dimensions remained consistent, with semantic dimensions maintaining a substantially larger influence (64.4%) on subjective identification than visual dimensions (28%).

These results suggest that while the overall predictability of subjective identification ratings diminishes post-disambiguation, the underlying reliance on semantic features remains robust. The minimal contribution of visual dimensions across conditions further reinforces the idea that subjective identification judgments are predominantly driven by higher-order semantic information, regardless of disambiguation.

### Semantic distance and entropy

Verbal accuracy of correct identification (Fig. 1c) provides valuable yet limited insights into the type of information gained during disambiguation. Here, we leveraged two complementary metrics to provide a more fine-grained characterisation of this process: semantic distance and semantic entropy. Semantic distance (see Fig. 3a) measures how far in the semantic space a provided label is concerning the target words or their synonyms. For instance, for a given image, participants may have typed a related but incorrect label (“donkey” for “horse”). A low semantic distance would then capture how closely related the answer is to the correct label. Quantifying these distances provides a more nuanced measure of how participants’ representations shift in semantic space after acquiring relevant information (i.e., during grey trials). Orthogonal to this measure, semantic entropy (see Fig. 3a) quantifies the degree of heterogeneity across participants in the labels provided for a given image. Lower semantic entropy reflects that participants are consistent in their responses to an image, using a small and constant set of labels. However, higher levels of semantic entropy indicate that participants use a broader set of labels when naming an object. A reduction in semantic entropy after being presented with the unambiguous grayscale image would suggest decreased variability in the labels provided by participants. Importantly, as mentioned before, the computation of these two metrics is independent (see Fig. 3a): a low semantic entropy does not imply a reduced semantic space, as a small set of responses (e.g. branch and tree; low entropy) can still be far from the target label (e.g. arm). For both metrics, our manipulation induced strong effects. For semantic distance (F_(2, 3616)_ = 3038, p < 0.001, η_p_^2^ = 0.63, 95% CI [0.61, 0.64]), the separation between the provided and true labels in the semantic space decreased after disambiguation (pre-disambiguation: M = 0.25, SD = 0.18; post-disambiguation: M = 0.46, SD = 0.22). Similarly, semantic entropy was significantly higher before than after disambiguation (pre-disambiguation: M = 2.52, SD = 0.92; post-disambiguation: M = 1.72, SD = 0.92; F_(2, 3616)_ = 2085, p < 0.001, ηp2 = 0.53, 95% CI [0.52, 0.55]). Finally, we tested the extent to which semantic distance tracks subjective identification. First, we labelled trials corresponding to individual images and participants based on their subjective identification pattern (e.g. “always identified” if a participant answered “yes” to the subjective identification question in all three versions of one image; pre, unambiguous grayscale, and post). We then assessed semantic distance for images following each of 4 a priori patterns: never identified, only identified during unambiguous grayscale trials, only identified in post and unambiguous grayscale trials (i.e. disambiguation pattern), and, finally, always identified. A pattern x exposure condition ANOVA showed a significant interaction of both factors (F_(6, 828)_ = 36.58, p < 0.001, η_p_ ^2^ = 0.21, 95% CI [0.16, 0.25]), revealing a differential effect of the condition on the semantic distance scores of images following different patterns of subjective identification (see Fig. 3b). More specifically, exposure had the greatest effect on semantic distance for those images only identified in their unambiguous version and post (t(138) = –19.82, p < 0.001, g = –1.97, 95% CI [-2.17, –1.76]), followed by images always identified (t = 11, p < 0.001), then images identified only in unambiguous grayscale (t(138) = –6.51, p < 0.001, g = –0.58, 95% CI [-0.76, –0.40]), and finally, images that were never identified (t(138) = –6.13, p < 0.001, g = –0.57, 95% CI [-0.76, –0.38]). Figure 3b also clarifies the relationship between subjective identification and semantic correctness. Trials in which participants reported identifying the object (‘always’ and ‘post-only’) showed substantially lower semantic distance than those in which they did not (‘never’), indicating closer alignment with the target concept. Notably, the ‘always’ group also showed lower semantic distance in the pre-disambiguation trial, suggesting that subjective identification reflects a perceptual judgement that precedes and partially overlaps with the accuracy of the verbal label. This distinction reinforces the idea that identification and naming capture different but related stages of the disambiguation process.

**Fig. 3.**
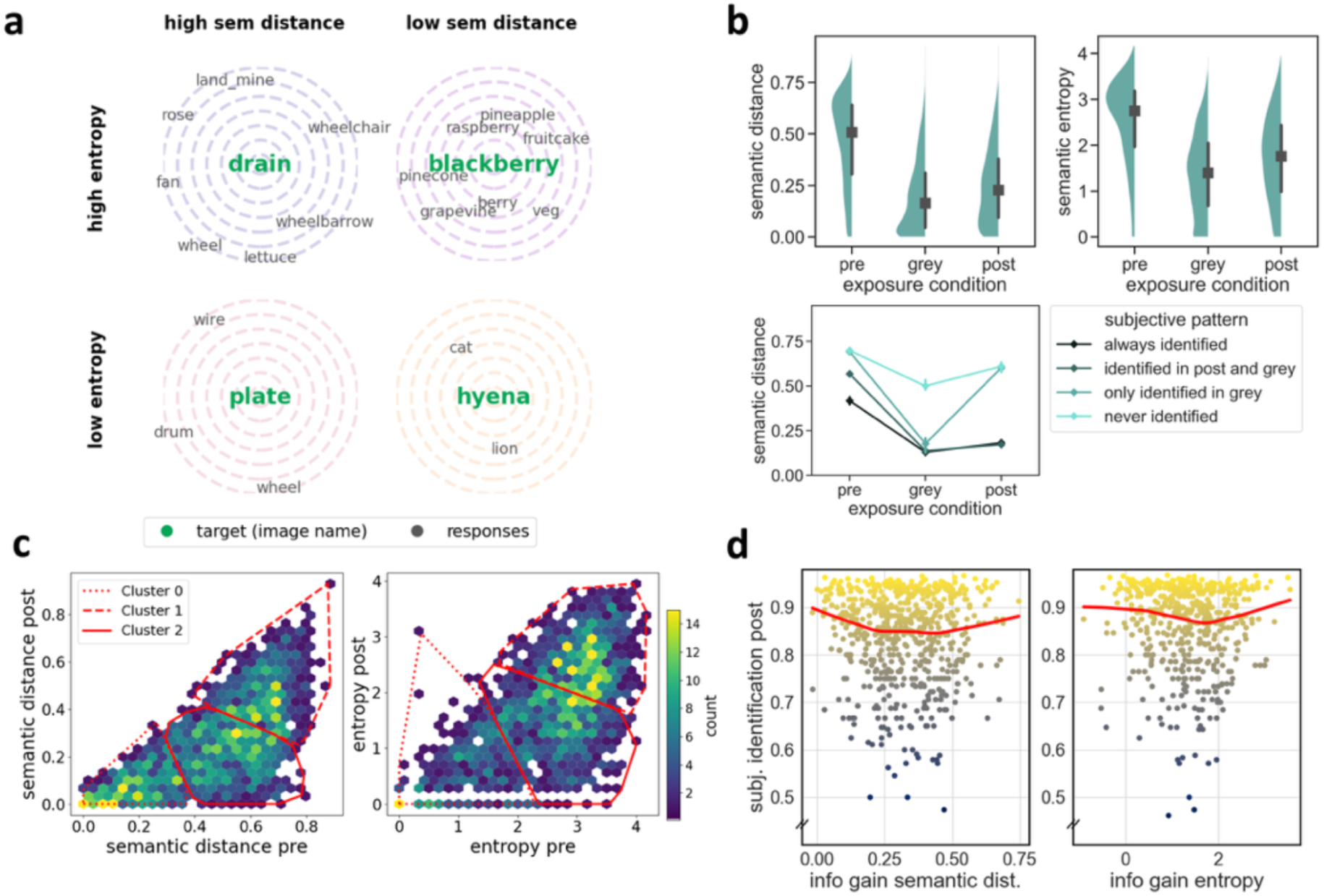
Semantic distance, entropy and their relation with subjective identification. **a,** Illustration of semantic distance and entropy. Targets (green dots) and responses (black dots) are shown in semantic space. High semantic distance indicates labels far from the target (e.g., “lettuce” for “drain”), while low semantic distance reflects closer labels (e.g., “raspberry” for “blackberry”). High entropy occurs when participants provide a broad range of labels, whereas low entropy reflects consistent responses. **b**, Effect of disambiguation on semantic measures. Top: Violin plots show reductions in semantic distance (left) and entropy (right) across exposure conditions (pre-, disambiguation [grey], and post-disambiguation). Black square represents medians, black lines denote interquartile range and green shaded areas indicate the data distribution. Bottom: Semantic distance across subjective identification patterns (always identified, never identified, only identified in disambiguation [grey], and only identified post) demonstrates the greatest effect of disambiguation on the images following the classic Mooney effect (“identified in post and grey” group). Markers indicate median semantic distance for each condition and error bars denote 95% confidence interval **c**, Hexbin plots display semantic distance (left) and entropy (right) from pre– to post-disambiguation. Red lines represent k-means clusters (k = 3), of which Cluster 2 shows stimuli of interest where values transition from high to low, reflecting significant information gain. **d**, Relationship between information gain and subjective recognition. Scatterplots show subjective identification post-disambiguation as a function of information gain in semantic distance (left) and entropy (right). Regression lines highlight a U-shaped relationship for semantic distance, suggesting that large initial gains reverse the trend at higher levels of semantic distance. For entropy, a similar but weaker quadratic pattern emerges. For illustrative purposes, data where subjective identifications were equal to 1 was excluded. The full set of images was used for the analyses reported in the main text. Semantic distance and entropy analyses were conducted on n = 1,809 images (five images excluded due to unavailable embeddings, see Methods). Regression analyses relating information gain to subjective identification were performed at the image level (n = 1,809 images), based on ratings from n = 947 participants.

In summary, these results demonstrate that the disambiguation of visual information leads to a reduced semantic distance and lower variability in participants’ responses. Altogether, this suggests that disambiguation not only sharpens semantic representation but increases the consistency in how visual information is interpreted across individuals.

### The complex relationship between information gained and later identification

One way of quantifying the amount of information acquired during disambiguation is computing the reduction in semantic distance and entropy between pre– and post-disambiguation trials of the same image (i.e. information gain). However, the impact of this gain on subsequent identification remains unclear. For instance, gaining more information could linearly improve identification (e.g., large increments of information would provide a richer context and thus lead to better identification performance than smaller increments). Yet, small increments of information can still entail profound changes in later identification albeit via different mechanisms (e.g., by confirming initial guesses ^31^). Notably, information gain in entropy and semantic distance were positively correlated, *r*(1807) = 0.655, *p* < 0.001, 95% CI [0.628, 0.681]. Hence, we employed a regression model (Ordinary Least Squares; OLS) to examine the effect of information gained either in semantic distance or entropy on subjective identification after disambiguation. To account for a non-linear (quadratic) relation between predictors and subjective identification, we also included squared terms for both independent variables.

When including all images, the model was statistically significant (F(4,1804) = 54.52, p < 0.001) and explained 10.8% of the variance in subjective identification (*R*² = 0.108, 95% CI [0.086, 0.132], adjusted *R*² = 0.106). Information gained in semantic distance and entropy showed a significant monotonic contribution for subjective identification, being such relation negative in the case of semantic distance (*t*(1804) = –10.907, *p* < 0.001, β = –0.7285, 95% CI [-0.8595, –0.5975]) and positive for entropy (*t*(1804) = 7.396, *p* < 0.001, β = 0.0695, 95% CI [0.0511, 0.0879]). Regarding a squared relationship between predictors and subjective identification, information gained on semantic distance had a significant non-linear effect (*t*(1804) = 6.185, *p* < 0.001, β = 0.7608, 95% CI [0.5196, 1.0021]) but not information gained in entropy (*t*(1804) = –0.414, *p* = 0.679, β = –0.0018, 95% CI [-0.0101, 0.0066]). We performed a split-half cross-validation analysis, fitting the regression model on one half of the data, and used it to predict identifiability in the other half across 1,000 iterations. The average correlation between predicted and observed identifiability closely resembled the original sample fit (*r* = 0.322, 95% CI [0.278, 0.363]), and the signs and relative magnitudes of the regression coefficients remained highly stable across splits. This pattern of results was also found when applying a jackknife approach, excluding data points where information gain in entropy or semantic distance fell below the 5th percentile or above the 95th percentile. In this trimmed dataset, the model remained significant (*F*(4, 1526) = 61.126, *p* < 0.001, *R*² = 0.138, adjusted *R*² = 0.136), and the overall pattern of results was preserved. Specifically, semantic distance showed a significant negative linear effect (*t*(1526) = –10.232, *p* < 0.001, β = –1.0623, 95% CI [-1.2659, –0.8586]) and a significant positive quadratic effect (*t*(1526) = 6.029, *p* < 0.001, β = 1.3179, 95% CI [0.8891, 1.7467]), while entropy also showed significant linear (*t*(1526) = 7.987, *p* < 0.001, β = 0.1467, 95% CI [0.1107, 0.1827]) and quadratic effects (*t*(1526) = –3.642, *p* < 0.001, β = –0.0346, 95% CI [-0.0532, –0.0160]). This confirms that the observed relationship is not merely driven by a small number of extreme values.

To specifically examine the role of gaining both types of semantic information after successful disambiguation, we focused this analysis on a subset of images exhibiting the most relevant pattern. Specifically, we selected images with high semantic distance and entropy values before disambiguation and lower values for both measures afterwards, corresponding to the bottom-right quadrant in both plots of Fig. 3c. To identify these images in a data-driven manner, we applied k-means clustering (k = 3) using pre– and post-disambiguation values for both variables (semantic distance and entropy). From this analysis, we selected Cluster 2 (see Fig. 3c), which displayed the desired pattern of interest: high values before disambiguation that dropped to lower values afterwards. Regression models were statistically significant for Cluster 2 both in semantic distance (*F*(2, 746) = 17.173, *p* < 0.001; *R*² = 0.044, adjusted *R*² = 0.041) and entropy (*F*(2, 705) = 12.271, *p* < 0.001; *R*² = 0.034, adjusted *R*² = 0.031). For semantic distance, results showed a significant negative linear relationship (*t*(746) = –5.602, *p* < 0.001, β = –0.4932, 95% CI [-0.6660, –0.3203]) and a positive quadratic relationship (*t*(746) = 4.895, *p* < 0.001, β = 0.6512, 95% CI [0.3900, 0.9124]). Similarly, for entropy, a significant negative linear relationship (*t*(705) = –4.514, *p* < 0.001, β = –0.0573, 95% CI [-0.0822, –0.0324]) and a positive quadratic relationship (*t*(705) = 3.478, *p* < 0.001, β = 0.0160, 95% CI [0.0070, 0.0251]) were observed. The same pattern of results was found when applying a jackknife approach for outliers (excluding data in information gained below or above the 5th and 95th percentiles, respectively), with significant linear and quadratic effects for both predictors, although effect sizes were attenuated in some cases. Together, these findings suggest that, while information gain initially relates negatively to subjective identification, a quadratic effect indicates that this relationship reverses at higher levels of semantic distance reduction and, to a lesser extent, entropy. In other words, small information gain might introduce uncertainty or challenge initial interpretation, which could lead to lower subjective identification. However, when information gain is low, confirming prior predictions, or large enough to violate previous expectations, it improves identification performance.

## Discussion

Despite the fact the world is inherently ambiguous, how humans deal with such ambiguity in natural vision remains unclear. What are the visual features underlying perceptual ambiguity and how do these influence its later resolution? How can we make use of new information to then shape the identification of stimuli that were perceptually uncertain? The present work provides insights into these questions by leveraging a combination of behavioural and computational modeling approaches to characterize ratings from over 1000 participants of a large-scale dataset of ambiguous Mooney images. In summary, we observed that (i) distortion of high– rather than low-level visual features induces initial ambiguity; (ii) after disambiguation, while low-level features increase their contribution to subjective identification, high-level ones decrease in relevance; (iii) disambiguation is characterised by information gain, i.e., a systematic reduction in semantic distance between ambiguous and unambiguous images, with a concurrent decrease in uncertainty; yet, (iv) the relationship between information gain and subjective identification follows a non-linear, U-shaped trajectory.

Previous research has explored the visual features that contribute to ambiguity, including spatial characteristics such as contours and edges, occlusion, and figure-ground relationships ^11^, as well as colour, luminance, motion, and depth. Our study extends previous attempts by capitalizing on DNNs that mimic the primate visual system to investigate how different visual features influence perception along the processing gradient of the ventral visual stream. Specifically, we examine this in a context where a series of low-pass filters are applied to unambiguous, clear images to generate Mooney versions of them (see Methods section for details). We found that in such manipulation, higher-level (vs. lower-level) features are particularly impaired. In other words, converting a clear image to a Mooney image mostly impairs higher-level visual features. But how does high-level visual information (or lack thereof) relate to resolving ambiguity? Pre-disambiguation (that is, when participants encounter a Mooney image for the first time before having seen its clear counterpart), the poorer preservation of high-level visual features in the Mooney image explains the observed low rate of identification. In contrast, after being able to reconstruct the semantic meaning of the Mooney image from the greyscale clear image (post-disambiguation phase), lower-level features become more strongly correlated with subjective identification. This indicates that lower-level features may play a more critical role for a successful match between the unambiguous and Mooney image versions. Therefore, even with identical sensory input, different representational content appears to be prioritized depending on the availability of a recently learned clarifying percept. Altogether, our results suggest that the resolution of visual ambiguity is not only a function of low-level visual feature properties but instead is highly dependent on transformations in higher-level visual components.

This transition from top-down guessing to bottom up matching could be interpreted from the perspective of the analysis-by-synthesis framework ^32^, in which visual perception alternates between generative prediction (synthesis) and evidence-driven verification (analysis). During pre-disambiguation, participants would stay in a synthesis mode, trying to identify the images based on their higher-level information. However, after disambiguation, participants may switch to an analysis mode, where identification is now mainly driven by matching the preserved low-level structure of the Mooney image to the internal representation acquired from the unambiguous version. This shift in strategy also aligns with predictive processing accounts of vision (e.g., ^2,4^). From this perspective, the unambiguous image would establish a high-precision prior (or top-down expectation) at the top of the hierarchy. Once this strong prior is in place, the system’s primary goal would shift from generating a hypothesis to “explaining away” the sensory prediction error at lower levels. As a consequence, the system would become more sensitive to how well the low-level input matches the detailed top-down prediction, effectively increasing the functional relevance of early visual areas.

Furthermore, this trajectory mirrors the Reverse Hierarchy Theory ^33^, which posits that while conscious perception often begins with a high-level gist (“vision at a glance”), detailed recognition requires a feedback process that re-recruits low-level areas (“vision with scrutiny”). Our results suggest that disambiguation acts as the catalyst for this re-entry, enabling the observer to access and utilize low-level feature information that was functionally inaccessible during the initial ambiguous state.

Together, our results therefore suggest that ambiguity resolution reflects a dynamic reorganisation of perceptual inference: from top-down hypothesis generation when sensory evidence is insufficient, to bottom-up template matching once the perceptual model is known.

Traditional perceptual learning metrics of ambiguity resolution (e.g., subjective identification, reaction times) have largely been agnostic to the type of information learned. Here, we found disambiguation to be characterised by a systematic reduction in semantic distance between an ambiguous image and its unambiguous greyscale version, with a concurrent decrease in entropy (i.e., less variance in the given labels for the same perceptual input). These two complementary metrics provide deeper insight into the information extracted from unambiguous images and its relationship with subjective identification during the post-disambiguation stage. Crucially, although both types of information track perceptual learning, the content captured by semantic distance and entropy differentially helps subjective identification. Across all trials, information gain in semantic entropy (i.e., reducing semantic entropy) relates positively to subjective identification, suggesting that reduced perceived ambiguity strengthens the subjective sense of identification. In contrast to this linear relationship, information gain in semantic distance relates positively to subjective identification only for high levels of information gain, with lower levels displaying a negative relationship. This distinction highlights the different roles played by these two metrics in perceptual learning: while semantic entropy tracks how well participants can perceive the object as an entity (e.g., “Now I clearly see the object”), semantic distance reflects the precision with which participants can access relational features of the stimuli (e.g., “The object is a horse-like animal”). The dataset that we are releasing with the current manuscript is thus well poised to further explore these metrics and their contributions to perceptual learning and other related phenomena.

Interestingly, when restricting analyses to images with reduced semantic distance/entropy after disambiguation (i.e., the frequently observed Mooney pattern), both metrics exhibit a non-linear relationship with subjective identification. Specifically, both minimal and maximal information gain in these metrics correspond to higher levels of subjective identification than moderate information gain. For semantic distance, maximal information gain occurs when the initial guess was semantically far from the correct label and was drastically reduced post-disambiguation. In contrast, minimal information gain describes trials where distance was similar during the initial guess and the post-disambiguation response. Whereas the former scenario arguably results in a large prediction error, which has been linked to knowledge updating ^34^, the latter one could reflect a situation where the initial guess was close enough to the actual stimulus and the clear image acted as a confirmation, enhancing subjective identification. Such duality finds parallels in accounts of the relationship between prediction error and episodic encoding ^35^, where the interplay of two opposing encoding mechanisms renders non-linear patterns in a common output variable ^31,36,37^. It is worth noting that these minimal-gain trials may reflect cases where participants had subjectively identified a percept (albeit with intermediate semantic distance) before disambiguation and maintained the same percept after disambiguation. In such cases, low semantic entropy may have additionally boosted the subjective feeling of identification and promoted a rigid internal model – that would only be updated in response to a substantial prediction error. Accordingly, only substantial reductions in semantic entropy lead to better subjective identification, again arguably via model updating. This speculative idea deserves future testing.

Altogether, the reported non-linear, U-shaped trajectory between information gain and subjective identification challenges the intuition that information gain linearly facilitates perception and suggests a more complex interplay where the dynamics between predictive mechanisms and prior knowledge must be taken into account. Relatedly, recent evidence has shown that post-disambiguation, ambiguous images elicit a neural representation more similar to its undistorted version than to the exact same image before disambiguation ^14,15^. This drastic representational change given the same sensory input suggests that encountering the distorted image after disambiguation brings back to mind the previously perceived clear version. This raises the interesting possibility that visual identification after gaining new information can be understood as a pattern completion process ^38^, in which a partial stimulus serves as a cue for a previously encoded unambiguous stimulus. However, to the best of our knowledge, the underlying processes behind the (re)activation of a partial (ambiguous) stimulus linked to prior non-ambiguous percepts have not been explored before. Although our results provide initial insights into this question, future research should ascertain the extent to which the association between ambiguous and unambiguous stimuli depends on connections between lower– and/or higher-level features.

In conclusion, we found that ambiguous, distorted stimuli primarily suffer from a loss of high-level visual features, while retaining lower-level features from their original, undistorted counterparts. Interestingly, while high-level information is initially crucial to subjective identification, lower-level features gain in relevance after successful disambiguation. This suggests that the successful reactivation of recently perceived prior knowledge relies more on matching low-level characteristics, whereas abstract and invariant object features play a greater role in identification before disambiguation. Finally, our results reveal a non-linear relationship between information gain and subsequent subjective identification. Altogether, these findings open future avenues for studying how information gain and predictive mechanisms interact in visual perception.

## Limitations

Although Mooney images offer a well-established and controlled approach to study visual disambiguation processes, they represent a specific form of visual ambiguity (two-tone and low-pass filtered images). Further research is needed to confirm whether the present findings can generalise to other sources of real-world ambiguity. Also, in this work we infer a perceptual and predictive mechanism from behavioural and computational modelling, without neural recording. For this reason, some claims regarding the underlying neural mechanisms (e.g., predictive processing, pattern completion or reactivation) are inferential and should be tested using neuroimaging techniques.

We also need to highlight that analyses on the preservation index and subjective identification are correlational and do not establish a direct causal role of particular feature levels in human disambiguation. Although our results reveal a shift in the contributions of different visual features, causal evidence for this shift would require converging neural or experimental manipulation approaches.

We also want to highlight some population and task constraints. First, although the study includes a large sample size, all participants were young, tested online and required to be native English speakers. These population constraints open important questions regarding the developmental and aging-related trajectories of the determinants of ambiguity resolution. Future studies should investigate these processes in children and older adults to determine the boundary conditions and generalizability of our findings. Second, although the task followed an established approach to study visual disambiguation across three phases (pre, disambiguation and post-disambiguation), the current study does not include cases where images were not explicitly disambiguated (i.e. catch trials; see for instance ^14^. This could limit a better understanding of the role of disambiguation trials compared to mere repetition of Mooney images.

## Data availability

The data that support the findings of this study are available at https://osf.io/mzp23/.

## Code availability

The code to reproduce the results of this study is available at https://doi.org/10.17605/OSF.IO/MZP23. All the materials plus a Mooney image database derived from the study can be found at https://github.com/wobc/things-mooney. This repository further contains the toolbox that was used by the experimenters to create the Mooney images as well as an interface to specify various criteria (e.g., semantic distances, recognition rates) and generate your own Mooney-style image set.

## Acknowledgments

J.L.D. was supported by Project PID2023-151104NA-I00 funded by MCIN/AEI/10.13039/501100011033 and by FEDER, EU, and Grant RYC2021-033940-I funded by MCIN/AEI/10.13039/501100011033 and by the European Union NextGeneration EU/PRTR. J.O.T. was supported by an European Research Council Starting Grant (ERC-2024-StG-CONNECTS-101161992), and Grant RYC2023-045452-I funded by MICIU/AEI /10.13039/501100011033 and by the FSE+. J.V. was supported by an FPI grant (CEX2021-001161-M-20-5) by the Spanish Ministry of Science and Innovation. M.H. was supported by a research group grant from the Max Planck Society, the ERC Starting Grant project COREDIM (ERC-StG-2021-101039712), and funding from the Hessian Ministry of Higher Education, Science, Research, and the Arts through a LOEWE Start Professorship and the Research Cluster “The Adaptive Mind” via its Excellence Program. C.G.G. was supported by Project PID2023-149428NB-I00 funded by MCIN/AEI/10.13039/501100011033 and by FEDER, EU, and Grant RYC2021-033536-I funded by MCIN/AEI/10.13039/501100011033 and by the European Union NextGeneration EU/PRTR. The Mind, Brain and Behavior Research Center receives funding from grants CEX2023-001312-M by MICIU/AEI/10.13039/501100011033 and UCE-PP2023-11 by the University of Granada. The funders had no role in the study design, data collection and analysis, decision to publish or preparation of the manuscript. This manuscript is part of the PhD thesis of J.V.

## Author contributions

J.L.D., J.O.T, and C.G.G. were responsible for conceptualization, data curation, formal analysis, funding acquisition, investigation, methodology, project administration, resources, software, visualization, and writing of the original draft and further edits. J.V. was involved in data curation, software, visualization and writing – review and editing. M.H. was involved in conceptualization, methodology and writing – review and editing.

