## Supplementary Material for "Determinants of visual ambiguity resolution"

#### Supplementary Methods

### Feature preservation for non-Money image transformations

To further contextualize our feature preservation approach, we conducted a control analysis using two additional image transformations: high- and low-pass spatial frequency filtering, being the latest an important step to create a Mooney-like image (see Methods section of the main manuscript for details). The main goal of this analysis was to assess whether the observed preservation profile in Mooney images was specific to this type of transformation and to what extent it was comparable to other types of more mainstream image transformations. In addition, this analysis allowed us to ascertain whether the use of similarity measures derived from CORnet-S representations could arbitrate between different types of transformations.

#### Image transformation

The 1854 images of the THINGS Plus database were transformed by removing either low-spatial frequencies (i.e., high-pass filter, created by subtracting a Gaussian low-pass filter with a sigma of 50 from the original frequency spectrum) or high-spatial frequencies (i.e., low-pass filter, created by applying a Gaussian filter with a sigma of 10 directly to the frequency spectrum). Thus, we created two new versions of each image: the high-pass filtered preserve fine details and edges while lacking coarse structures; in contrast, the low-pass filtered lacks fine details but preserves broader structures. The filtering was performed in the frequency domain using a Fourier transform, followed by a multiplication with the respective Gaussian mask, and then reconstructed via inverse Fourier transform.

#### Feature preservation computation

We followed the same procedure used for the Mooney transformation (see Main text for more details). In short, we ran both high-pass and low-pass versions of each image through CORnet-S and extracted feature representations at each hierarchical layer (V1, V2, V4, IT) using the Net2Brain toolbox. To quantify feature preservation, we computed the Pearson correlation between the feature sets of each transformed image and its corresponding greyscale counterpart at each layer. This yielded preservation indices for each high-pass and low-pass transformed image.

### Results

The preservation index was significantly higher than zero ( Fig. 2b) across all layers after both transformations (for low-pass transformation, V1: $t(1853) = 518.523$, $p < 0.001$, Cohen's D = 12.042, V2: $t(1853) = 382.815$, $p < 0.001$, Cohen's D = 8.891, V4: $t(1853) = 369.324$, $p < .001$, Cohen's D = 8.577, IT: $t(1853) = 134.965$, $p < 0.001$, Cohen's D = 3.134; for high-pass transformation, V1: $t(1853) = 126.653$, $p < 0.001$, Cohen's D = 2.941, V2: $t(1853) = 366.811$, $p < 0.001$, Cohen's D = 8.519, V4: $t(1853) = 371.891$, $p < 0.001$, , Cohen's D = 8.637, IT: $t(1853) = 127.867$, $p < 0.001$, Cohen's D = 2.970). For low-pass filtered images, the mean preservation index was higher for V1 (M = 0.867, SD = 0.072) and decreased in V2 (M = 0.616, SD = 0.069), V4 (M = 0.560, SD = 0.065), and IT (M = 0.340, SD = 0.108). A one-way ANOVA was conducted to examine whether the preservation index differed across layers, revealing a statistical significant difference, $F(3,7412) = 13368$, $p < 0.001$, $\eta p2 = 0.844$, (CI for V1: [0.864, 0.870], CI for V2: [0.613, 0.619], CI for V4: [0.557, 0.563], CI for IT: [0.335, 0.345]), indicating that, similar as with the Mooney transformation, the original features of the greyscale image were particularly impaired in higher-level layers with the low-pass filter. In contrast to the low-pass filter, for high-pass filtered images, the mean preservation index was roughly equivalent across all layers (V1: M = 0.477, SD = 0.162; V2: M = 0.566, SD = 0.066; V4: M = 0.602, SD = 0.070; IT: M = 0.388, SD = 0.131). However, a one-way ANOVA conducted in the same way as above still revealed a main effect of layer, $F(3,7412) = 1292$, $p < 0.001$, $\eta p2 = 0.343$ (CI for V1: [0.470, 0.484], CI for V2: [0.563, 0.569], CI for V4: [0.599, 0.605], CI for IT: [0.382, 0.394]).

.

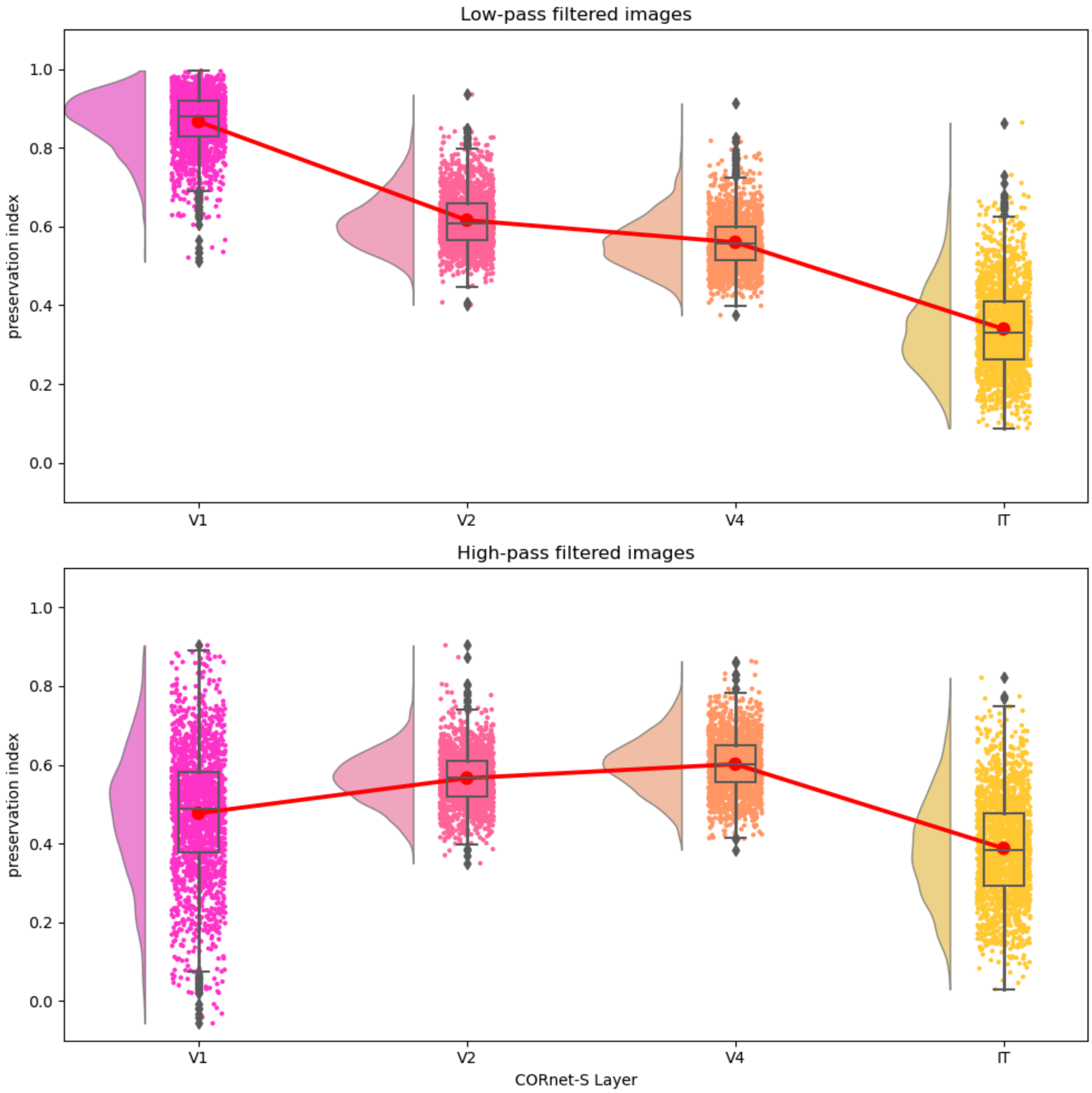


**Supplementary Figure 1. Preservation indices across CORnet-S layers for filtered images.** *Preservation indices for low-pass filter transformation (upper panel) and for high-pass filtered transformation (lower panel). Similar to the Mooney transformation, preservation in the low-pass filtered images decreases progressively from early visual layers (V1) to higher-level layers (IT), indicating that higher-level representations are more impaired. In contrast, high-pass filtered images display a comparable preservation index across layers, reflecting a strong impairment of features represented in early visual layers.*

Further analysis of how the preservation index changed across layers revealed significant differences in slopes between transformations ($F(2,5553) = 5669.64$, $p < 0.001$, CI for Mooney: [-0.1515, -0.1492], CI for low-pass: [-0.1653, -0.1620], CI for high-pass: [-0.0260, -0.0203]). Slopes were calculated using linear regression (stats.linregress) with layers encoded numerically (V1=0, V2=1, V4=2, IT=3) as the independent variable and Preservation Index as the dependent variable for each concept within each transformation type. For the Mooney transformation, there was a strong negative slope ($M = -0.164$, $SD = 0.036$), indicating a consistent decrease in preservation across successive layers. The low-pass filtered images showed a similar pattern with a negative slope ($M = -0.150$, $SD = 0.024$), suggesting that both transformations led to progressive degradation of the original image features at higher layers. In contrast, high-pass filtered images showed a smaller negative slope ($M = -0.023$, $SD = 0.063$), consistent with the more uniform preservation observed across layers. Post-hoc comparisons using Bonferroni-corrected t-tests revealed significant differences between all transformation pairs. The slope for high-pass filtered images was significantly less negative than both Mooney ($t(3700) = -80.76$, $p < 0.001$, Cohen's D: -2.655, CI: [-0.1303, -0.1241]) and low-pass filtered images ($t(3706) = -83.06$, $p < .001$, Cohen's D: -2.728, CI: [0.0114, 0.0153]), while Mooney images showed a slightly but significantly less negative slope than low-pass filtered images ($t(3700) = 13.17$, $p < 0.001$, Cohen's D: 0.433, CI: [0.1372, 0.1439]). These results suggest that while all transformations led to some degradation of feature preservation in higher layers, this effect was markedly stronger for low-pass filtered and Mooney images compared to high-pass filtered images, where feature preservation remained more stable across the processing hierarchy.

The similar pattern between the Mooney and the low-pass filter transformation is consistent with the fact that the process of creating the Mooney images included a low-pass filtering stage. Finally, the distinct preservation profiles (i.e, the significantly different slopes across transformations) validate our approach as a sensitive measure capable of capturing meaningful differences in how visual features are preserved across different layers for different types of image transformations.
